# Octahedral small virus-like particles of dengue virus type 2

**DOI:** 10.1101/2024.10.17.618815

**Authors:** Adam Johnson, Martín Dodes Traian, Richard M. Walsh, Simon Jenni, Stephen C. Harrison

## Abstract

Flavivirus envelope (E) and precursor M (prM) proteins, when ectopically expressed, assemble into empty, virus-like particles (VLPs). Cleavage of prM to M and loss of the pr fragment converts the VLPs from immature to mature particles, mimicking a similar maturation of authentic virions. Most of the VLPs obtained by prM-E expression are smaller than virions; early, low-resolution cryo-EM studies suggested a simple, 60-subunit, icosahedral organization. We describe here the cryo-EM structure of immature, small VLPs from dengue virus type 2 and show that they have octahedral rather than icosahedral symmetry. The asymmetric unit of the octahedral particle is an asymmetric trimer of prM-E heterodimers, just as it is on icosahedral immature virions; the full, octahedrally symmetric particle thus has 24 such asymmetric trimers, or 72 prM-E heterodimers in all. Cleavage of prM and release of pr generates ovoid, somewhat irregular, mature particles. Previous work has shown that mature smVLPs have fusion properties identical to those of virions, consistent with local, virion-like clustering of 36 E dimers on their surface. The cryo-EM structure and the properties of these VLPs described here relate directly to on-going efforts to use them as vaccine immunogens.

**IMPORTANCE:** Ectopic expression of flavivirus envelope (E) and precursor M (prM) proteins leads to formation and secretion of empty, virus-like particles (VLPs), which are candidate, non-infectious, virion-like components of flavivirus vaccines. We show that the immature particles of a major class of VLPs -- “small VLPs” (smVLPs), which have smaller diameter than those of virion size, -- are octahedrally (rather than icosahedrally) symmetric, with the same clustering of prM and E, as asymmetric trimers of prM-E heterodimers, found on immature virions. Cleavage of prM and formation of mature, smVLPs yields somewhat irregular, ovoid particles. Design and characterization of VLPs as vaccine components will need to take these properties into account.

## INTRODUCTION

Flaviviruses assemble by budding into the endoplasmic reticulum (ER) as immature particles, pass through the secretory pathway, and emerge from the cell as mature, infectious virions. The icosahedrally symmetric, immature particles contain 180 envelope (E) subunits associated with the same number of membrane-protein precursor (prM) subunits and internal, core (C) protein subunits (Fig. 1A). Both E and prM are glycoproteins with C-terminal, transmembrane anchors (Fig. 1B). The conserved E-protein ectodomains (DI, DII, and DIII) connect to the helical-hairpin anchors through a segment traditionally called the “stem” (a term that antedates any structure). The C subunits, on the cytosolic side of the ER membrane, interact with the 10.7 kb, plus-sense, RNA genome and incorporate it into the particle as it assembles and buds into the ER lumen.

**FIG 1.**
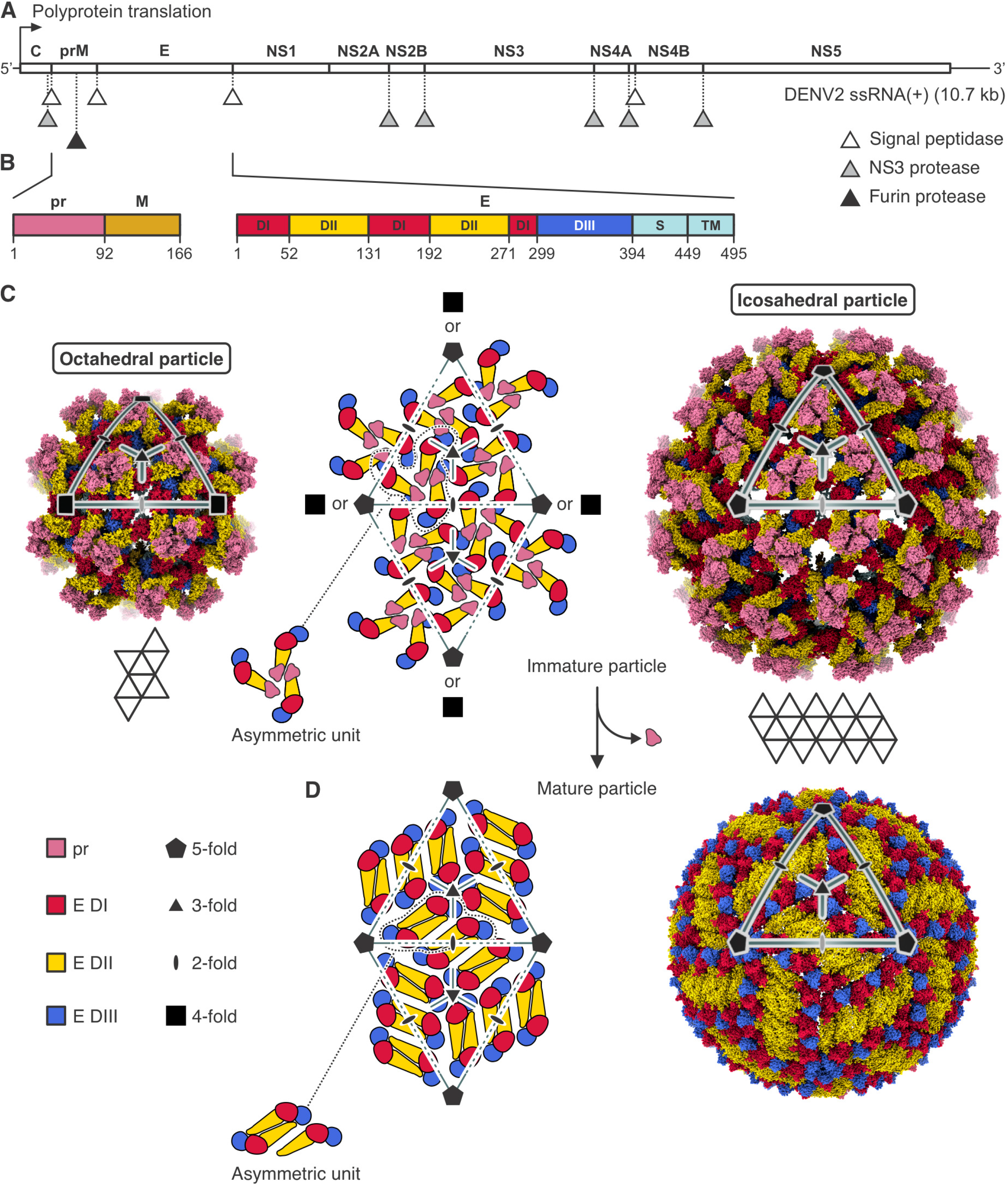
Flavivirus structural organization. (A) Flavivirus proteins are produced in the host cell from an approximately 10.7 kb long single-stranded positive sense genome/messenger RNA, ssRNA(+). A polyprotein is translated, which is co- and post-translationally processed by different host and viral proteases (signal peptidase, NS3 protease, furin protease) at positions indicated by triangles. C, (anchored) capsid protein; prM, membrane glycoprotein precursor; E, envelope protein; NS1, nonstructural protein 1; NS2A, nonstructural protein 2A; NS2B, nonstructural protein 2B; NS3, nonstructural protein 3 (protease); NS4A, nonstructural protein 4A; NS4B, nonstructural protein 4B; NS5, nonstructural protein 5 (methyltransferase and RNA-dependent RNA polymerase). (B) Primary sequence and domain organization of the prM and E proteins. Residues at domain boundaries are numbered. pr sequence is colored pink, M is brown. E protein domains are labeled I (red), II (yellow), III (blue), S (stem, cyan), and TM (transmembrane region, cyan). prM is cleaved by furin protease after residue 91. (C) Structural organization of immature flavivirus particles. The middle panel shows schematically the arrangement of prM-E protomers on a pseudohexagonal lattice. Folding of 20 triangular faces around 5-fold axes results in icosahedral particles, as the one shown in the right panel, which is the structure of an infectious virion (immature SPOV particle, PDB-ID 6ZQW). Folding of 8 triangular faces around 4-fold axes results in octahedral particles, as the one shown in the left panel, which is the structure of the DENV2 smVLP reported here. In both structures, the octahedral and icosahedral particles, the triangulation number is T = 1 with 3 prM-E protomers per asymmetric unit. Protein domains are colored as in (B). (D) Structural organization of mature flavivirus particles. Proteolytic cleavage by furin in the TGN, exposure to mildly acidic pH, and release of virions from the host cell with dissociation of the pr protein, is accompanied by substantial structural rearrangements of the M and E proteins to generate the mature particle, in which M and E form dimers. The structure of the mature particle has the same icosahedral symmetry as the immature particle (mature SPOV particle, PDB-ID 6ZQV).

Sixty, markedly asymmetric clusters of three prM-E protomers cover the surface of an immature particle (Fig. 1C, Fig. S1 and S2) (1-6). Ninety, twofold symmetric M-E dimers cover the surface of a mature particle (Fig. 1D, Fig. S1 and S2) (7, 8). The large-scale reorganization of the particle surface, from immature to mature, probably occurs in the trans-Golgi network (TGN), in response to the mildly acidic pH of the TGN lumen (3). The reorganization exposes a furin site on prM; cleavage at that site will allow release of the N-terminal, “pr” fragment when the particle again reaches neutral pH as it emerges from the infected cell. During infection of a new target cell, exposure to low pH in an endosome triggers a further fusogenic rearrangement of E into post-fusion trimers (9).

Cryo-EM structures have been determined for several mature flavivirus particles, but only two for immature virions at high resolution (5, 6) and one at lower resolution (10). The structure of immature Binjari virus (BinJV) has defined the previously uncertain connectivity between the pr fragment, which covers the fusion loop of E, and the membrane-associated C-terminal fragment (which becomes “M” in the mature particle) (5). While it is likely that the connectivity in BinJV is generally true, as inferred from sequence conservation (Data set S1–S3 in the supplemental material), density in the structure of Spondweni virus (SPOV) is too poorly defined in the connecting segments to confirm it (6).

Ectopic expression of flavivirus prM-E alone leads to production and secretion of virus-like particles (VLPs), which undergo the same immature-to-mature transition as virions and which have virion-like fusion properties (11-13). Particles of two or more sizes result. Particles in one class closely resemble virions in size (about 500 Å in diameter) and isometric shape; those in a second class (smVLPs) are smaller, with about one-third the number of E protein subunits on their surface (14, 15). The latter predominate in our preparations (13, 16) and in several of those reported in the literature (15). Early work suggested that the smVLPs of tick-borne encephalitis virus (TBEV) were 60-subunit, icosahedral particles (17). But because icosahedral symmetry for a 60-subunit surface requires symmetric three-fold clusters, it is incompatible with the asymmetric threefold clusters now known (from work that came several years later) to be present on immature virions and presumably on the smVLPs. We show here that immature smVLPs of dengue virus type 2 (DENV2), and by extension, those of other flaviviruses, are octahedrally symmetric particles, with prM-E asymmetrically clustered as on immature virions. The octahedral asymmetric unit is one asymmetric cluster of three prM-E heterodimers, and thus there are 72 prM-E heterodimers in total. The transition to mature particles, generated by cleavage in vitro, produces somewhat ovoid particles, presumably covered by 36 E dimers.

## RESULTS AND DISCUSSION

We obtained a 3D reconstruction of an immature DENV2 smVLP at 6.5 Å resolution (Fig. 2A and B, Fig. S5A). Focused classification and refinement of a symmetry-expanded particle image stack yielded an improved map of the asymmetric unit (3 prM-E heterodimers) at 4.3 Å resolution (Fig. 2C, Fig. S5B). We obtained an AlphaFold 2 structure from the DENV2 prM-E sequence, docked 3 models of the prM-E heterodimer independently into the 3 subunits of the asymmetric unit, and manually modeled the linker sequence between pr and M. The model was fit to the locally focused map by domain-wise rigid-body fitting, structure-morphing and real space refinement (Fig. 2C, S5B, S5C). The stem and transmembrane segments of E and prM were reasonably well defined in the map, in particular for the two of the three protomers (Fig. 3). The sharper curvature of the particle displaced the transmembrane segments from their positions, relative to their ectodomains, in the 180-subunit, immature flavivirions for which subnanometer structures are known (Fig. 3). Our map is consistent with the revised prM connectivity seen clearly in the structure of immature BinJV (5), in which the three pr domains at the tips of an asymmetric threefold cluster connect directly to the three stem and transmembrane anchors (stem-TMs) clustered around the same pseudo-threefold (Fig. 3A). In our locally refined map, essentially continuous density (weak from residues 215-224, but unambiguously traced for two of the chains and suggested by very weak density for the third) connects each “pr” domain associated with a fusion-loop tip of E with a corresponding stem-TM. One local difference is in the orientation of the prM stem-TM associated with the E subunit marked B in Fig. 3B. In our octahedral smVLP structure, the segment that connects the pr domain and stem-TM of the prM in question allows the B-subunit stem-TM to form an approximately twofold symmetric contact with the stem-TM of prM associated with subunit C -- that is, to orient the stem-TM about 180° from its orientation in the full particle (Fig. 3B, C). Moreover, this contact is essentially the same as the twofold contact between M proteins in the mature icosahedral particle. As suggested in early work on dengue virus particles and confirmed by the BinJV structure, the furin site is inaccessible in the immature particle but becomes exposed on the virion surface during the immature-to-mature lattice transition (Fig. 3B).

**FIG 2.**
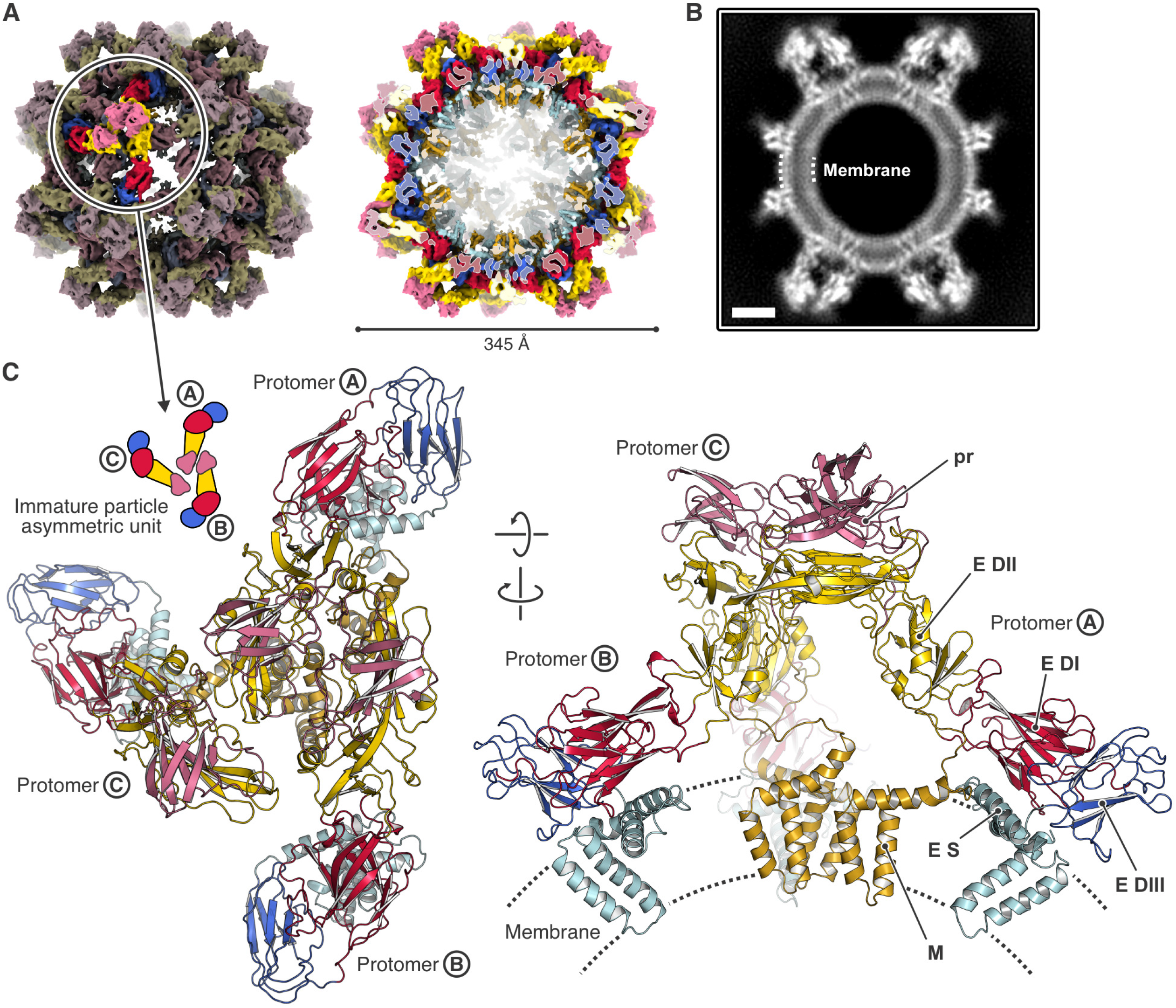
Structure of the dengue virus type 2 (DENV2) octahedral small virus-like particle (smVLP). (A) cryo-EM map of the octahedral smVLP at 6.5 Å resolution, viewed along the 2-fold symmetry axis. prM-E protein domains are colored according to the following scheme: pr, pink; M, brown; E DI, red; E DII, yellow; E DIII, blue; E stem and C-terminal domains, cyan. Left, one asymmetric unit, consisting of an asymmetric trimer of three prM-E protomers (labeled A, B, and C), is highlighted. Right, the particle is cut to allow inside view of the M and E transmembrane α helices. (B) Projection of a 12 Å-thick central slice of the octahedral smVLP cryo-EM map showing density arising from the lipid bilayer. The scale bar corresponds to 50 Å. (C) Ribbon representation of the asymmetric timer structure, viewed form the outside of the particle (left), and viewed from the side (right). The membrane is schematically indicated. Domains are colored as in (A).

**FIG 3.**
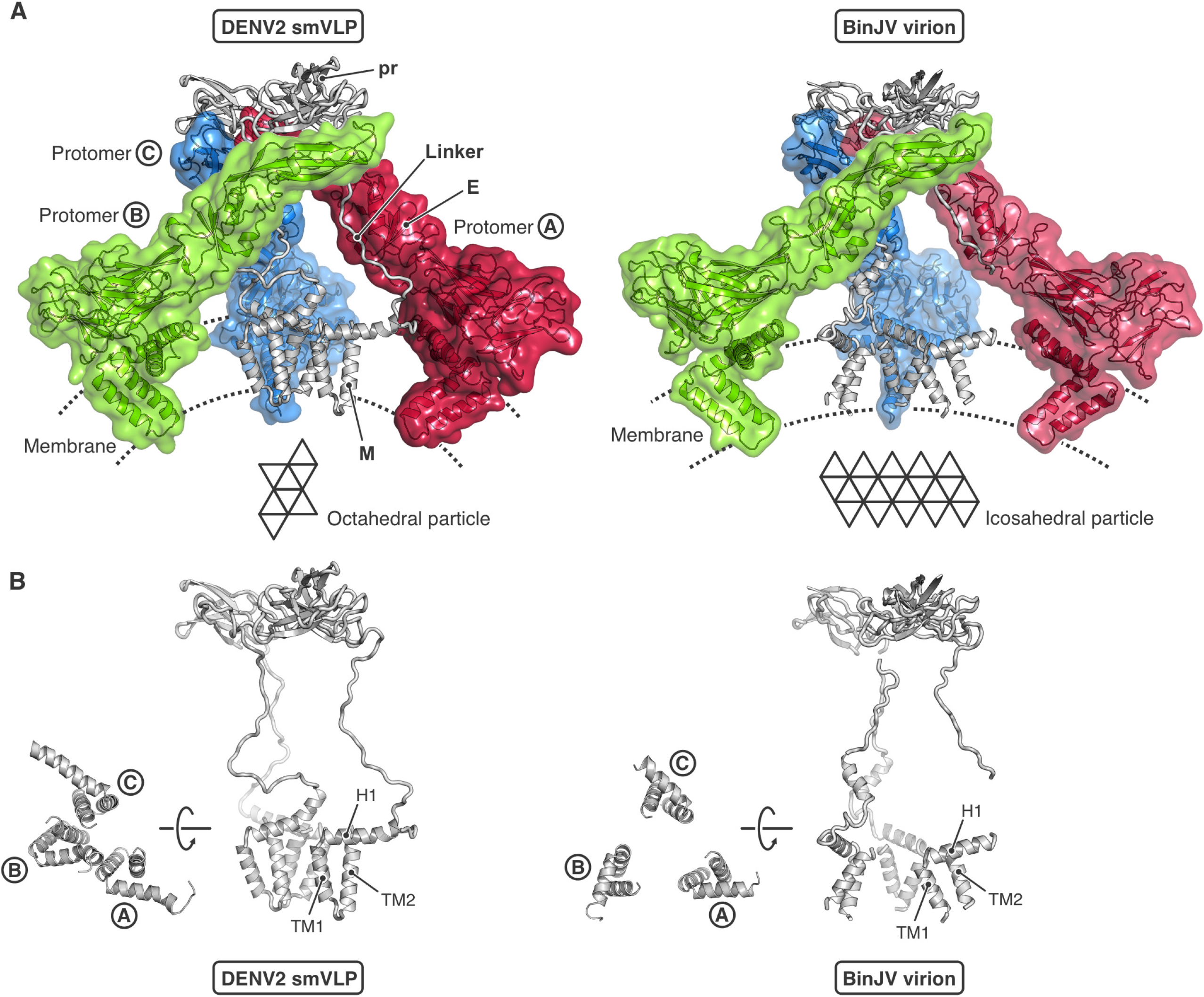
Comparison of the prM-E asymmetric trimer from the octahedral DENV2 smVLP and the icosahedral BinJV immature flavivirus virion. (A) DENV2 smVLP (left) and BinJV virion (right, PDB-ID 7L30). The E subunits are shown in transparent surface representation and colored differently: protomer A, red; protomer B, green; protomer C, blue. prM is shown in gray ribbon representation for all three protomers. The octahedral DENV2 smVLP (left) contains 72 prM-E protomers, with 3 asymmetric trimers within each of the 8 triangular faces (3 × 3 × 8 = 72). The icosahedral immature BinJV virion (right) contains 180 prM-E protomers, with 3 asymmetric trimers within each of the 20 triangular faces (3 × 3 × 20 = 180). (B) View of the prM within one asymmetric trimer for the octahedral DENV2 smVLP (left) and the icosahedral immature BinJV virion (right).

How does the subunit packing on the immature particle surface transform into the arrangement on the surface of a mature particle? The pattern of rearrangement proposed for BinJV (5) suggests (i) that the E subunits marked B and C in Fig. 4A rearrange into the BC dimer, with the C subunits clustered around the fivefold (fourfold in case of our smVLP), and (ii) that one of the three E subunits related by the threefold axis (one of which is marked A) pair to form an AA’ dimer (across an icosahedral twofold). As the authors of that paper point out, this model requires a non-icosahedrally symmetric intermediate. But symmetry breaking would then require that the transition propagate asymmetrically from the proposed starting point. Moreover, broken symmetry seems hard to reconcile with the reversibility of the transition under in vitro conditions in which exposure of the furin site does not lead to its cleavage (18), as also pointed out by the authors of the SPOV structure (6). An alternative would be to pair the two A subunits related by an icosahedral twofold in the immature form (A’ and A” in Fig. 4A, B), requiring those subunits to move and rotate substantially, but allowing retention of icosahedral symmetry throughout the transition and symmetrically related pairings across the entire the particle. The SPOV paper illustrates this alternative in its supplementary movie 2 (6). In principle, it would be possible to rotate B and C’ in the opposite direction, so that B subunits cluster on the fivefold in the mature structure. Either alternative is compatible with retention of icosahedral symmetry. Subunits B and C’ are related to each other by a local twofold in both the immature and mature lattices (asterisks in Figs. 4A and 4B) and also in the octahedrally symmetric, immature smVLP (asterisk in Fig. 2A). Thus, BC’ pairing retains local twofold symmetry. Retention of this local twofold would also retain interactions in both the immature and mature lattices between the stem-TM of prM and the stem-TM of E, because the stem-TM of the E subunit marked B contacts the stem-TM of prM associated with the subunit marked C’, and vice versa.

**FIG 4.**
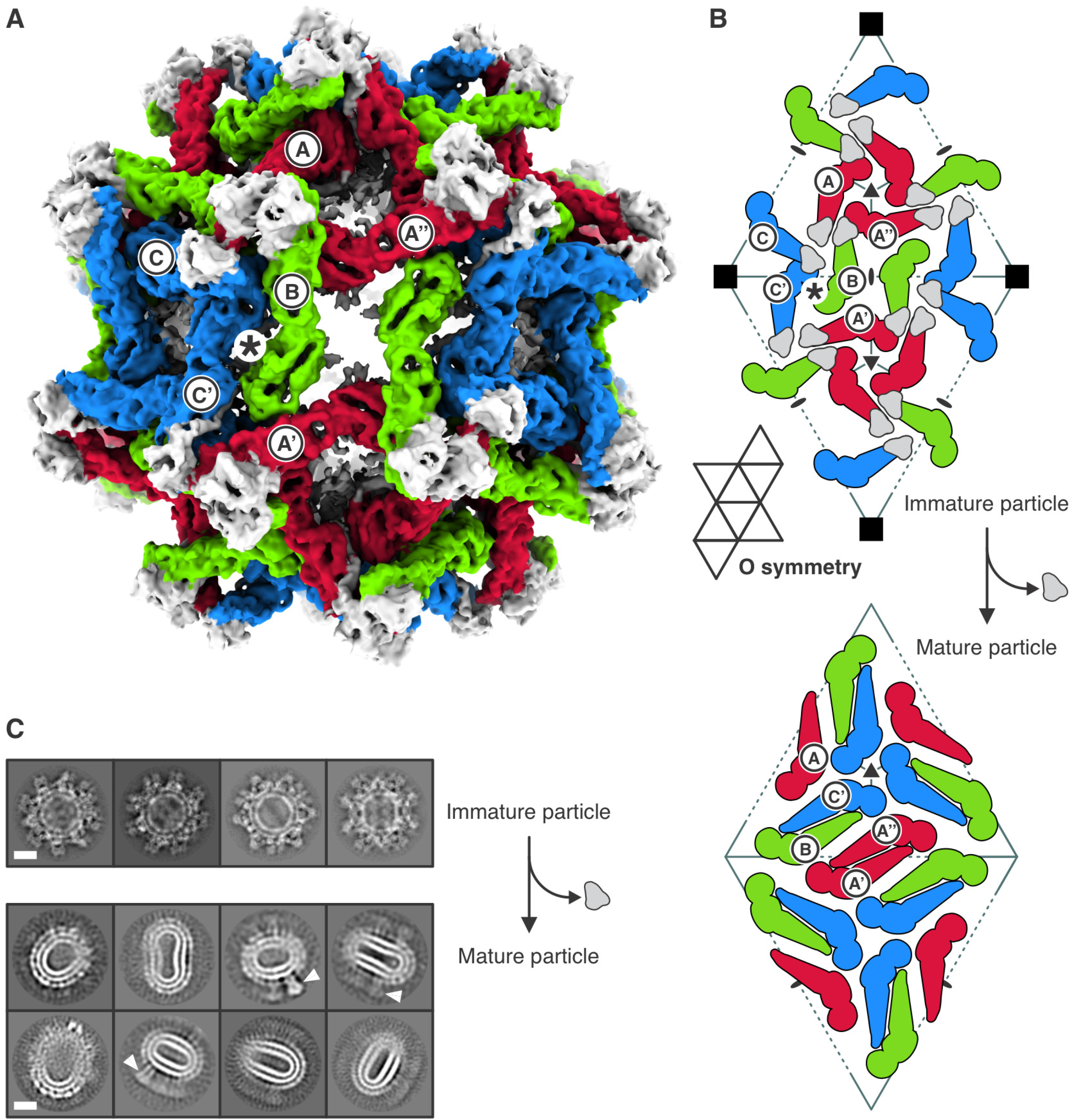
Immature and mature conformation of DENV2 smVLPs. (A) Structure of the octahedral particle with the asymmetric trimers in immature conformation. Protomers are colored red, green, and blue. prM of all protomers is shown in gray. Protomers of the reference asymmetric unit are labeled A, B, and C. Protomers of neighboring asymmetric units are labeled with a prime or double prime, e.g. A’, and A’’. The position of a pseudo 2-fold axis relating protomers B and C’ is indicated by an asterisk. (B) Arrangement of prM-E protomers in the immature particle (top). Were octahedral symmetry retained during maturation (cleavage of pr-M and loss of pr), rearrangement would yield the packing of M-E protomers shown (bottom). The elongated, mature smVLPs show, however, that octahedral symmetry is not retained. (C) 2D class averages of immature (top) and mature (bottom) DENV2 smVLPs. White arrowheads indicate what appear to be E trimers in post-fusion conformation. The scale bar corresponds to 100 Å.

The fusion properties of mature smVLPs of tick-borne encephalitis virus (TBEV) (12) and West Nile virus (WNV) (13) are indistinguishable from those of virions. Moreover, the kinetics of fusion by dengue smVLPs (all four serotypes) closely resemble those of WNV smVLPs (16). Cleavage of prM and release of the pr fragment must therefore allow E to rearrange as dimers on the smVLP surface into a packing that resembles, at least locally, the organization of E dimers on a virus particle. Otherwise, we would expect the kinetics and pH-dependence of fusion to be different. Our mature dengue smVLPs are too irregular for a cryo-EM reconstruction (Fig. 4C), but studies of TBEV smVLPs, before the asymmetry of the immature trimer was known, suggested a T=1 icosahedral structure (17). A similar interpretation has also been described for cryo-EM reconstructions of mature dengue type 2 smVLPs (15). How can we reconcile those observations with a 72-subunit immature precursor? We regard it as unlikely that subunits (and lipids) could be lost during a transition. The analysis of TBEV smVLPs was based on selected particles that were circular in outline, and they could therefore have been ovoid particles viewed near or along their major axis. The same might have been true of the dengue smVLPs examined by (15), in which particles were selected from a heterogeneous mixture of sizes. Clustering of BC dimers around a five-fold axis is a principal feature of subunit packing on the surface of mature virions, and it is therefore plausible that a few, similar five-fold interactions would form as the smVLPs convert from immature to mature (see Fig. 4B and description in caption). Imposition of icosahedral symmetry in calculating the reconstruction would inevitably have yielded a symmetric map, probably with low-resolution features of the local fivefold interactions we suggest will form.

VLPs are candidates for development of safe, flavivirus vaccines (15, 19, 20). Depending on the virus, expression strategies and production conditions, the particles are probably a mixture of small and full-size VLPs. Epitope exposure is likely to be somewhat different on the mature versions of the two particle sizes, and vaccine trials with such particles will need to pay attention to the mixture present in the immunogen.

## MATERIALS AND METHODS

### Purification of DV2 VLPs

We produced DV2 VLPs as described (13, 16), with some modifications. HEK 293T cells stably expressing the DV2 prME gene were grown as adherent cultures by passaging them in DMEM High Glucose medium supplemented with 10% FBS, 1% PenStrep, and 150 μg/ml puromycin. On day 1, cells were transferred into suspension culture at 1 million cells/ml in 293 Freestyle media and incubated for 4 days at 28 °C. Cells were then removed by centrifugation and VLPs precipitated from the supernatant by adding PEG-8000 and NaCl to final concentrations of 8% (w/v) and 0.5 M, respectively, and incubating overnight at 4 °C. The mixture was then centrifuged at 3500 g, and the pellet resuspended in 20 mM HEPES pH 7.5, 150 mM NaCl for 20 min at 4 °C. Centrifugation at 5000 g for 20 min clarified the solution, and the supernatant t was then applied to a 5/10/15/20/60% (w/v) sucrose step gradient and centrifuged for 20 h in a SW41 rotor at 38,000 rpm. The 20% (w/v) sucrose fraction was collected and buffer exchanged to 20 mM HEPES pH 7.5, 150 mM NaCl with an Amicon 100 kDa cutoff concentrator. VLPs were concentrated to 1 mg/ml protein as estimated by SDS-PAGE and Coomassie staining.

### Cryo-EM grid preparation and data collection

Samples were vitrified using a Cryoplunge 3 robot. We applied 3.5 μl purified VLPs (1 mg/ml total protein concentration) to copper mesh holey carbon grids (C-flat 1.2 μm diameter holes with 1.3 μm spacing and a support thickness of ∼20 nm), blotted for 4 s and flash-frozen in liquid nitrogen-cooled liquid ethane. The humidity was kept at >90% during freezing. We collected movie image stacks with a Gatan K2 Summit direct detector on a TF30 Polara electron microscope operating at 300 kV accelerating voltage using SerialEM (21). The calibrated magnification at the physical pixel size was 40650 and the defocus range approximately between -1.0 and -3.0 μm, resulting in a physical pixel size of 1.23 Å at the image plane. We collected three datasets in super-resolution counting mode with an electron dose rate of 8 electrons per physical pixel area per second and an exposure time of 200 ms per frame (40 frames per movie), resulting in a total exposure time of 8 s and a total electron dose of 42.3 electrons per Å^2^ per movie (1.06 electrons per Å^2^ per frame).

### Cryo-EM data processing

Movie frames were aligned (5 × 5 patch), averaged and binned two times to a physical pixel size of 1.23 Å with MotionCor2 (22). Initial contrast transfer function (CTF) parameters were determined with CTFFIND4 (23) from the summed micrographs. We first calculated a low resolution smVLP map with octahedral symmetry imposed from a subset of manually picked particles using Relion (24), which was then used to generate reference projections with EMAN2 (25) for particle picking with Gautomatch (Fig. S3A). After local defocus estimation with Gctf (26) and 2D classification in Relion (24), we retained 38,934 particle images with a box size of 512 × 512 pixels (Fig. S3B). Particles were aligned with cisTEM (27) (refine3d version 1.01, reconstruct3d version 1.02), where we imposed octahedral symmetry (O) and limited the alignment resolution to 8 Å. After local movie frame alignment and determination of optimal weighting factors for frame summation (relion_motion_refine), and estimation of CTF aberrations (relion_ctf_refine) (28), the nominal resolution of the octahedral smVLP reconstruction was 6.5 Å as determined by the Fourier shell correlation (FSC) between half maps (Fig. S4A and Table S1), similar to what was previously reported for cryo-EM reconstructions of SPOV (6) and BinJV (5) immature virus particles. For a local reconstruction, focusing on a single smVLP asymmetric unit, we symmetry expanded the particle stack, signal-subtracted the 6.5 Å-resolution density of the smVLP (except for the single asymmetric unit in question), and extracted a 256 × 256-pixel particle stack containing 934,416 images, each centered at one of the of the symmetry-related positions. After alignment by classification without alignment (29), the images and associated metadata were imported into CryoSPARC (30). A 3D classification into 10 classes without alignment was applied to the image stack; particles that partitioned into lower resolution classes were discarded. This step led to a stack of 706,684 particles and a reconstruction at 4.49 Å resolution. Another round of 3D classification into 4 classes led to a stack of 403,357 particles, from which a final local refinement was performed with adaptive marginalization and non-uniform refinement (31), limiting the rotational search to 20° and the shifts to 10 Å. The final reconstruction had a nominal resolution of 4.24 Å (Fig. S4B and Table S1). Data analysis, modeling, and refinement software was curated by SBGrid (32).

### Structure modeling and refinement

We used AlphaFold 2 (33) to obtain a model of a prM-E protomer, for which we input the sequences for pr, M, and E, respectively, as individual chains. From the resulting model, we placed the following domains into the corresponding densities of the three protomers in the 4.3 Å-resolution map of the local reconstruction: pr (residues 1–91), M (residues 110–166), E DI (residues 1–51, 131–191, 271–298), E DII (residues 52–130, 192–270), and E DIII (residues 299– 393). Part of the E-protein stem and C-terminal domain (residues 394–449 out of residues 394– 495) was taken from the model of the immature SPOV particle (PDB-ID 6ZQJ) and mutated to the DENV2 sequence (Data set S1 in the supplemental material), because AlphaFold 2 modeled the structure in the mature conformation and therefore this part did not fit the density of the immature particle. Domains were rigid-body fit and adjusted to the density with phenix.real_space refine using morphing (34). We then used RosettaCM (35) to fix the connections between the fitted domains. 3- and 9-mer fragment libraries were obtained from the Robetta server (http://robetta.bakerlab.org). We incorporated secondary structure restraint terms and the fit to the density map in the RosettaCM scoring function (36). We used the program O (37) to manually model the linker sequences connecting the pr and M domains (residues 92–109). The complete model consisting of prM-E protomers A, B, and C was refined with phenix.real_space (34). To obtain a model of the octahedral particle, we took the refined model of the asymmetric unit, rigid-body fit it into the 6.5 A-resolution octahedral map, and symmetry-expanded the structure to generate the full particle. The FSC between the refined models and the cryo-EM maps are shown in Fig. S4C and D. Map and model statistics are summarized in Table S1.

### Sequence alignments

Flavivirus sequences (Data set S1–S3 in the supplemental material) were retrieved from the NCBI Viral Genomes Resource (38) with the following accession codes: dengue virus type 2 (DENV2), NP_056776.2; dengue virus type 1 (DENV1), NP_059433.1; dengue virus type 3 (DENV3), YP_001621843.1; dengue virus type 4 (DENV4), NP_073286.1; Japanese encephalitis virus (JEV), NP_059434.1; West Nile virus (WNV), YP_001527877.1; St. Louis encephalitis virus (SLEV), YP_001008348.1; Spondweni virus (SPOV), YP_009222008.1; Zika virus (ZIKV), YP_002790881.1; Powassan virus (POWV), NP_620099.1; Tick-borne encephalitis virus (TBEV), NP_043135.1; Yellow fever virus (YFV), NP_041726.1. Sequences were aligned with MAFFT (39) and multiple sequence alignment were printed and annotated with ESPript (40).

### Figure preparation

Figures were prepared using PyMOL (The PyMOL Molecular Graphics System, Version 2.3 Schrödinger, LLC), ChimeraX (41), matplotlib (42) and ImageMagick (ImageMagick Studio LLC, 2023, available at: https://imagemagick.org).

## Data availability

Cryo-EM maps and refined models have been deposited in the Electron Microscopy Data Bank and Protein Data Bank, respectively, with accession identifiers EMD-47082 and PDB-ID 9DOF for the local reconstruction of the asymmetric unit, and EMD-47083 and PDB-ID 9DOG for the full octahedral smVLP.

## ACKNOWLEDGMENTS

We thank Zongli Li for help with electron microscopy.

## FIGURE CAPTIONS

**FIG S1** Schematic illustration of subunit packing in immature and mature flavivirus particles with a domain coloring scheme (domains I, II, III in red, yellow and blue, respectively). The folding of a hypothetical planar, hexagonal (p6) lattice follows the scheme of Caspar and Klug (43). ((A) The immature conformation of prM-E on a p6 lattice with an asymmetric trimer within each asymmetric unit. (B) Folding of triangular faces to generate the 5-fold axes of an icosahedral immature particle with one asymmetric trimer within the icosahedral asymmetric unit (T=1). Each sixfold axis of the p6 lattice becomes a fivefold axis of the icosahedral particle. (C) Folding of triangular faces to generate the 4-fold axes of an octahedral immature particle. Each sixfold axis of the p6 lattice becomes a fourfold axis of the octahedral particle. (D) The mature conformation of M-E on a p6 lattice with one and a half dimers within each asymmetric unit. (E) Folding of triangular faces to generate the 5-fold axes of an icosahedral mature particle.

**FIG S2** Schematic illustration of subunit packing in immature and mature flavivirus particles with a protomer coloring scheme. Protomers or each asymmetric unit are colored red, green, and blue. prM of all protomers is shown in gray. The folding of a hypothetical planar, hexagonal (p6) lattice follows the scheme of Caspar and Klug (43). (A) The immature conformation of prM-E on a p6 lattice with an asymmetric trimer within each asymmetric unit. (B) Folding of triangular faces to generate the 5-fold axes of an icosahedral immature particle. (C) Folding of triangular faces to generate the 4-fold axes of an octahedral immature particle. (D) The mature conformation of M-E on a pseudo hexagonal lattice with one and a half dimers within each asymmetric unit. (E) Folding of triangular faces to generate the 5-fold axes of an icosahedral mature particle.

**FIG S3** cryo-EM analysis of DENV2 smVLPs. (A) Low pass filtered micrograph. Particles picked and retained for processing are circled. The scale bar corresponds to 500 Å. (B) 2D class averages of immature DENV2 smVLPs. The scale bar corresponds to 500 Å.

**FIG S4** Fourier shell correlation (FSC) analysis. (A) and (B) FSC between half maps for the octahedral and local reconstructions, respectively. The nominal resolution at which the correlation drops below 0.143 is shown. Light blue, unmasked; dark blue, after applying a soft mask to the half maps. (C) and (D) FSC between the final map and the model for the octahedral and local reconstructions, respectively. The nominal resolution at which the correlation drops below 0.5 is shown. Light red, unmasked; dark red, after applying a soft mask to the final map.

**FIG S5** Local resolution and *B* factor analysis. (A) cryo-EM reconstruction of the full DENV2 smVLP with octahedral symmetry. (B) Local reconstruction of the asymmetric trimer. (C) *B* factors mapped on the refined structure of the asymmetric trimer. *B* factor values are only meaningful relative to other values in the structure, as they depend on the degree of sharpening that was applied to the cryo-EM reconstruction.

**Data set S1** Multiple sequence alignment of part of the flavivirus polyprotein covering prM-E. DENV2, dengue virus type 2; DENV1, dengue virus type 1; DENV3 dengue virus type 3; DENV4, dengue virus type 4; JEV, Japanese encephalitis virus; WNV, West Nile virus; SLEV, St. Louis encephalitis virus; SPOV, Spondweni virus; ZIKV, Zika virus; POWV, Powassan virus; TBEV, tick-borne encephalitis virus; YFV, yellow fever virus. Secondary structure elements are indicated above the sequences. Solid bars below the sequences are colored according to the following domains: pr, pink; M, brown; E DI, red; E DII, yellow; E DIII, blue; E stem and C-terminal domains, cyan. Proteolytic cleavage sites are indicated by double ovals. See Materials and Methods for NCBI Viral Genomes Resource accession codes.

**Data set S2** Flavivirus prM protein multiple sequence alignment. DENV2, dengue virus type 2; DENV1, dengue virus type 1; DENV3 dengue virus type 3; DENV4, dengue virus type 4; JEV, Japanese encephalitis virus; WNV, West Nile virus; SLEV, St. Louis encephalitis virus; SPOV, Spondweni virus; ZIKV, Zika virus; POWV, Powassan virus; TBEV, tick-borne encephalitis virus; YFV, yellow fever virus. Secondary structure elements are indicated above the sequences. Solid bars below the sequences are colored according to the following domains: pr, pink; M, brown. The furin proteolytic cleavage sites is indicated by double ovals. See Materials and Methods for NCBI Viral Genomes Resource accession codes.

**Data set S3** Flavivirus E protein multiple sequence alignment. DENV2, dengue virus type 2; DENV1, dengue virus type 1; DENV3 dengue virus type 3; DENV4, dengue virus type 4; JEV, Japanese encephalitis virus; WNV, West Nile virus; SLEV, St. Louis encephalitis virus; SPOV, Spondweni virus; ZIKV, Zika virus; POWV, Powassan virus; TBEV, tick-borne encephalitis virus; YFV, yellow fever virus. Secondary structure elements are indicated above the sequences. Solid bars below the sequences are colored according to the following domains: E DI, red; E DII, yellow; E DIII, blue; E stem and C-terminal domains, cyan. See Materials and Methods for NCBI Viral Genomes Resource accession codes.

---

**TABLE S1.** Cryo-EM data collection and model statistics.

| Octahedral small virus-like particles of dengue virus type 2 |  |  |
| --- | --- | --- |
| **Data collection** |  |  |
| Electron microscope | Polara |  |
| Magnification | 40,650 |  |
| Voltage (kV) | 300 |  |
| Defocus range (μm)<sup>a</sup> | 1.5–3.0 |  |
| Physical pixel size (Å) | 1.23 |  |
| Number of movies | 13,808 |  |
| **Octahedral reconstruction** |  |  |
| Number of images | 38,934 |  |
| Box size (pixels) | 512 |  |
| Symmetry imposed | O |  |
| Map resolution (Å)<sup>b</sup> | 6.5 |  |
| **Local reconstruction** |  |  |
| Number of images | 403,357 |  |
| Box size (pixels) | 256 |  |
| Symmetry imposed | C<sub>1</sub> |  |
| Map resolution (Å)<sup>b</sup> | 4.3 |  |
| **Model statistics** | **Local reconstruction** | **Octahedral reconstruction** |
| EMD accession identifier | EMD-47082 | EMD-47083 |
| PDB accession identifier | 9DOF | 9DOG |
| Refinement resolution (Å) | 4.3 | 6.5 (rigid body fitting only) |
| CC (mask) | 0.67 | 0.71 |
| Model composition |  |  |
| Non-hydrogen atoms | 15,354 |  |
| Protein residues | 1,983 |  |
| *B* factors (Å<sup>2</sup>) |  |  |
| Min | 65 |  |
| Max | 782 |  |
| Mean | 174 |  |
| R.m.s. deviations |  |  |
| Bond lengths (Å) | 0.005 |  |
| Bond angles (°) | 0.948 |  |
| Validation |  |  |
| MolProbity clash score | 8.4 |  |
| Poor rotamers (%) | 0.0 |  |
| Ramachandran plot |  |  |
| Favored (%) | 92.6 |  |
| Allowed (%) | 7.1 |  |
| Disallowed (%) | 0.3 |  |

<sup>a</sup> Approximate range of underfocus.<sup>b</sup> Resolution where FSC between masked half maps drops below 0.143.

